# Integrative multiomics analysis of metabolic dysregulation induced by occupational benzene exposure in mice

**DOI:** 10.1101/2024.12.22.629805

**Authors:** Sydney Scofield, Lisa Koshko, Lukas Stilgenbauer, Alix Booms, Roxanne Berube, Christopher Kassotis, Chung-Ho Lin, Hyejeong Jang, Seongho Kim, Paul Stemmer, Adelheid Lempradl, Marianna Sadagurski

## Abstract

**Background:** Type 2 Diabetes Mellitus (T2DM) is a significant public health burden. Emerging evidence links volatile organic compounds (VOCs), such as benzene to endocrine disruption and metabolic dysfunction. However, the effects of chronic environmentally relevant VOC exposures on metabolic health are still emerging.

**Objective:** Building on our previous findings that benzene exposure at smoking levels (50 ppm) induces metabolic impairments in male mice, we investigated the effects of occupationally relevant, below OSHA approved, benzene exposure on metabolic health.

**Methods:** Adult male C57BL/6 mice were exposed to 0.9ppm benzene 8 hours a day for 9 weeks. We assessed measures of metabolic homeostasis and conducted RNA and proteome sequencing on insulin-sensitive organs (liver, skeletal muscle, adipose tissue).

**Results:** This low-dose exposure caused significant metabolic disruptions, including hyperglycemia, hyperinsulinemia, and insulin resistance. Transcriptomic analysis of liver, skeletal muscle, and adipose tissue identified key changes in metabolic and immune pathways especially in liver. Proteomic analysis of the liver revealed mitochondrial dysfunction as a shared feature, with disruptions in oxidative phosphorylation, mitophagy, and immune activation. Comparative analysis with high-dose (50 ppm) exposure showed both conserved and dose-specific transcriptomic changes in liver, particularly in metabolic and immune responses.

**Conclusions:** Our study is the first to comprehensively assess the impacts of occupational benzene exposure on metabolic health, highlighting mitochondrial dysfunction as a central mechanism and the dose-dependent molecular pathways in insulin-sensitive organs driving benzene-induced metabolic imbalance. Our data indicate that current OSHA occupational exposure limits for benzene are insufficient, as they could result in adverse metabolic health in exposed workers, particularly men, following chronic exposure.

## Introduction

Exposure to air pollutants in the environment is a globally rising health risk [1]. Recent studies show that exposure to volatile organic compounds (VOCs), a prominent subgroup of environmental pollutants, is associated with obesity, dyslipidemia, liver injury, and increased risk of Type 2 Diabetes Mellitus (T2DM) [2, 3]. Moreover, a recent meta-analysis established a robust association between exposure to benzene, a prevalent airborne VOC, and insulin resistance in humans across all ages [2, 4]. While exposure can occur in ambient indoor and outdoor environments, an important source of exposure to consider is occupational, where pollutant levels may be at much higher concentrations than in the regular environment [5]. Evidence shows that there are substantial VOC toxicants present in the workplace across the globe, and exposure to these toxicants represents a significant risk factor in the development of disease [6–10].

In the United States, the Occupational Health and Safety Administration (OSHA) regulates the maximum average exposure to airborne benzene during an 8-hour workday as 1 ppm [11]. Given that workers may encounter these benzene levels daily, it is crucial to investigate potential long-term health effects. Previously, we showed that exposure to benzene at concentrations found in cigarette smoke induces metabolic dysfunction and neuroinflammation in adult male mice, with microglial inflammation playing a key role [4, 12]. Here, we aimed to understand how chronic exposure to benzene at sub-permitted regulatory levels (0.9 ppm) affects whole-body metabolism and compared these effects to a higher smoking-level exposure. We now demonstrate that exposure to benzene at levels below OSHA approved occupational threshold is sufficient to cause metabolic impairments similar to those observed at higher doses.

Additionally, transcriptional profiling of insulin-sensitive tissues revealed both shared and unique gene signatures driving these metabolic effects. Our study demonstrates a critical gap in regulatory guidelines, suggesting that currently approved occupational exposure levels do not account for the impact on metabolic health and predisposition to diabetes over time. Therefore, our study is one of the first to comprehensively assess the metabolic dysfunction arising from occupationally relevant benzene doses in mice, thus contributing to the broader implications of benzene exposure on human metabolic health.

## Materials and Methods

### Animals

9-week-old C57BL/6 male mice were obtained from The Jackson Laboratory. Male mice were used exclusively in this study since females showed no benzene-induced metabolic effects as reported previously [12]. Animals were kept on a 12-hour light cycle and had access to food and water *ad libitum*. Mice were exposed to benzene (0.9 ppm) during the light cycle for 8 hr/day, 7 days/week, for 9 weeks (**Fig. 1A**). All exposures were performed using the FlexStream automated Perm Tube System (KIN-TEK Analytical, Inc) as previously described [12]. Tissues were collected following measurements of energy expenditure. All animal procedures were approved by Wayne State University IACUC and performed in accordance with the National Institutes of Health Guide for the Care and Use of Laboratory Animals.

**Figure 1.**
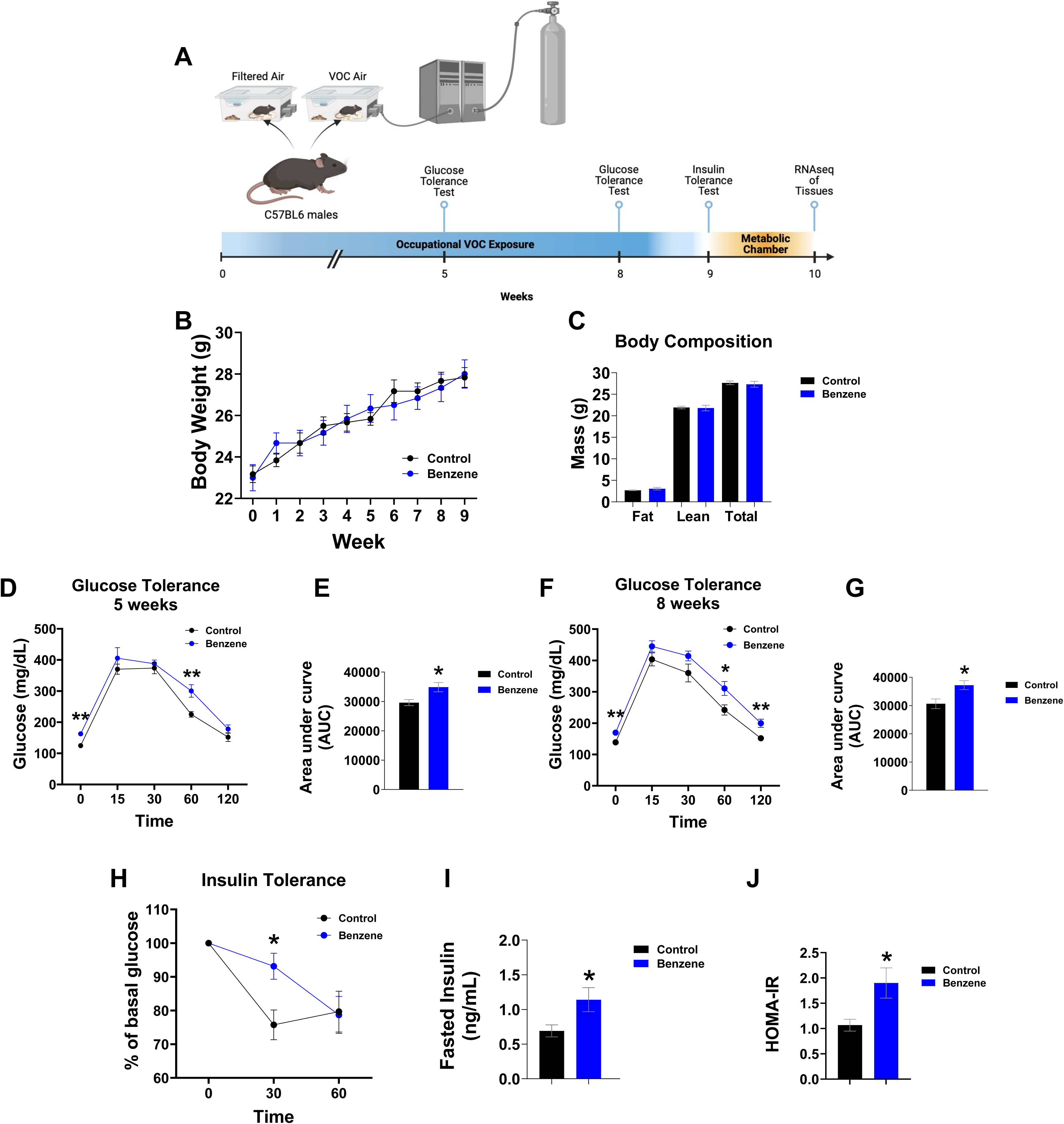
Prolonged occupational benzene exposure disrupts glucose and insulin tolerance in adult male mice. (A) Experimental timeline. (B) Body weight and (C) body composition. (D) Glucose tolerance test and (E) area under the curve following 5 weeks of exposure. (F) Glucose tolerance test and (G) area under curve following 8 weeks of exposure. (H) Insulin tolerance test after 9 weeks of exposure. (I) Fasted insulin levels after 8 weeks of exposure. (J) Homeostatic Assessment of Insulin Resistance (HOMA-IR). Data shown as the mean ± SEM (n=6/group). Repeated measures analysis of variance followed by Newman-Keuls post-hoc test, or t-test. (*,p<0.05, **, p<0.01).

### Metabolic measurements

For glucose tolerance test (GTT), mice were fasted for 6 hr and intraperitoneally injected with D-glucose (Sigma-Aldrich Cat # 158968-1KG) at a dose of 2 g/kg⋅BW. Blood glucose levels were measured using a Contour Next EZ meter at basal state (0 min) and then at 15, 30, 60, and 120 minutes after injection as before [13]. For insulin tolerance test (ITT), mice were fasted for 4 hours followed by intraperitoneal injection of Humulin R (Humalog NDC # 0002-7510-01) (0.8U/kg body weight). Blood glucose was measured at 0, 30, and 60 minutes. Blood insulin was determined on serum from tail vein bleeds using Insulin ELISA kit (Crystal Chem.Inc. #90080). Parameters of energy expenditure (VO_2_ consumption, VCO_2_ production, respiratory exchange ratio (RER) and heat production were measured by indirect calorimetry using the Bruker Phenomaster (TSE Systems, Germany #160407-03).

During this time, animals were single-housed and kept on a 12-hour light cycle with access to food and water *ad libitum*. After a 24-hour acclimatization, data were collected and used for analyses.

### Urinary tt-MA levels

Following the exposure, urine was collected and immediately frozen at −80° C until further processing. 20uL of the urine sample were transferred into a vial containing 800uL of 100% methanol. The sample was fortified with isotopic labeled trans,trans-Muconic-d4 acid as the internal standard (IS). The t,t-Muconic acid (t,t-MA) and the was isotopic labeled IS were extracted by vortexing for 2 min. Following the extraction, the extract was filtered with a 0.2 um Whatman Anotop filter. The concentrations of t,t-MA and IS were determined by a Waters Acquity Ultra-High-Performance Liquid Chromatography (Waters, Milford, MA, USA) coupled with a XEVO TQ-XS tandem mass spectrometer (UHPLC-MS/MS, Waters) controlled by MassLynx software Ver 4.2. The analytes were separated by a CORTECS® C18+ analytical column with 1.6 µm particle size, 100 mm length x 2.1 mm internal diameter connected to CORTECS® UPLC C18 VanGuard Pre-Column (1.6 µm particle size, 5 mm length x 2.1 mm internal diameter). Separation was achieved using a linear gradient of 0.01% formic acid in water (A) and 0.01% formic acid in 100% acetonitrile (B). The gradient conditions are: 0–0.2 min, 2% B; 0.2–1.89 min, 2–80% (linear gradient) B; 1.89-1.92 min, 80–98% (linear gradient) B; 1.92–3.61 min (linear gradient), 98% B; 3.61–6.77 min, 2% B with a flow rate of 0.4 ml/min. The column temperature will be set at 40 C and the autosampler temperature will be set at 10 C. The system will be first conditioned with 50 % acetonitrile and 50% of 0.01% formic acid, and the column will be equilibrated with 2% acetonitrile and 98% of 0.01% formic acid solution before injection. The injection volume is 1 µl. The mass analyzer Xevo-TQXS equipped with an electrospray ionization source will be operated in negative ion mode (ES-). The acquisition parameters for t,t-MA and trans,trans-Muconic-d4 acid (IS) were carried out in the multi-reaction monitoring mode (MRM) with the deprotonated precursor ion [M − H]−. The fragment ions monitored: m/z 140.95➔52.96 and m/z 144.91➔56.99 for quantification of analytes t,t-MA and IS, respectively. The ionization energy, multi-reaction monitoring (MRM) transition ions (precursor and product ions), capillary and cone voltage (CV), gas flow, and collision energy (CE) were optimized by Waters IntelliStart™ optimization software package. The data was processed, quantified, and reviewed by TargetLynx software Ver 4.2. The optimized cone ionization energy was 6V with optimized collision energy 10eV for t,t-MA and 8eV for IS trans,trans-Muconic-d4 acid. Urinary tt-MA levels were normalized to creatinine levels. Urinary creatinine was measured by colorimetric assay (Crystal Chem. Cat # 501947694).

### RNA extraction

For RNA extraction, tissue samples were stored in RNA-later (Invitrogen, #AM7020) at −80°C. The tissues were handled on dry ice, weighed, and immediately frozen with liquid nitrogen in screw cap tubes containing 3 mm Zirconium Beads (OPS Diagnostics, #BAWZ 3000-300-23). 1 mL TRIzol Reagent (Life Technologies, #15596026) was added per 50-100 mg of tissue. Tubes were homogenized using the FastPrep-24 system on high for 30 seconds and visually observed for tissue disruption, performing additional homogenizations as needed. Samples were vortexed for 15 seconds and incubated for 5 minutes at room temperature. 200 ul chloroform (Sigma-Aldrich, #319988) was added, followed by vortexing for 15 seconds and incubation at room temperature for 2 minutes. Samples were centrifuged at 12,000 x g for 15 minutes at 4°C. The upper aqueous phase was transferred to new Eppendorf tubes followed by RNA precipitation using 500 ul ice-cold isopropanol (Sigma-Aldrich, #190764) and 2 ul GlycoBlue (ThermoFisher, #AM9516) for pellet visualization. Samples were incubated for 10 minutes at room temperature, then centrifuged at 12,000 x g for 15 minutes at 4°C. Supernatant was removed and discarded. The samples were washed twice with 75% ethanol (Fisher Scientific, #111000200), using a 5-minute 7,500 x g centrifugation each round for pellet collection. The pellet was air-dried for 10 minutes and resuspended in 20 ul RNase-free water (Life Technologies, #AM9938). Samples were incubated at 60°C for 5 minutes, then stored at −80°C.

### Construction and sequencing of RNA-seq libraries

Libraries were prepared by the Van Andel Genomics Core from 500 ng of total RNA using the KAPA RNA HyperPrep Kit (Kapa Biosystems, Wilmington, MA USA). Ribosomal RNA material was reduced using the QIAseq FastSelect –rRNA HMR Kit (Qiagen, Germantown, MD, USA). RNA was sheared to 300-400 bp. Before PCR amplification, cDNA fragments were ligated to IDT for Illumina TruSeq UD Indexed adapters (Illumina Inc, San Diego CA, USA). Quality and quantity of the finished libraries were assessed using a combination of Agilent DNA High Sensitivity chip (Agilent Technologies, Inc.), QuantiFluor® dsDNA System (Promega Corp., Madison, WI, USA), and Kapa Illumina Library Quantification qPCR assays (Kapa Biosystems). Individually indexed libraries were pooled and 50 bp, paired-end sequencing was performed on an Illumina NovaSeq6000 sequencer to an average depth of 30M raw paired reads per transcriptome. Base calling was done by Illumina RTA3 and the output of NCS was demultiplexed and converted to FastQ format with Illumina Bcl2fastq v1.9.0.

### Data analysis

Adapter and low-quality sequences were trimmed from the reads using Trim Galore v0.6.0 with the ‘--paired’ parameter. Trimmed reads were aligned to the mm10 genome with GENCODE M24 gene annotations[14], using STAR v2.7.8a [15] with the parameters ‘--twopassMode Basic’, and ‘--quantMode GeneCounts’ to generate the read counts per gene. Downstream analyses were conducted in iDEP 2.01 [16] using the reverse-stranded counts produced by STAR. Transcripts with a total raw read count < 0.5 counts per million (CPM) were filtered out from the analysis. Principal component analysis (PCA) plots for liver and muscle RNA-seq are shown in **Figure S2A, B**. Read counts were analyzed using DEseq2 in iDEP 2.01 to detect differentially expressed genes (DEGs) using FDR cutoff of 0.1 and minimum fold change of 1.5. DEGs were further processed for pathway analysis using iDEP enrichment analysis for the enrichment of KEGG pathways, Reactome, WikiPathways, and gene ontology (GO) terms (biological processes, molecular function, and cellular component) with a FDR cutoff of 0.05. Plots were created with SR Plot (bioinformatics.com.cn/srplot). Cluster diagrams were created using Cytoscape ClueGO [17]. RNA-Seq analysis for adipose tissues was performed at the WSU Genome Sciences Core as before (Debarba et al. 2024). All RNA-Seq data are available at the Sequence Read Archive (SRA) at NCBI #PRJNA1194550.

### Protein extraction for proteomic analysis

Liver tissue was homogenized using a Storm 24 Bullet Blender (Next Advance) with 100 ul of unbuffered water and 3.2 mm stainless steel beads. Following the first round of homogenization 100 ul of 5% Lithium dodecyl sulfate detergent (LiDS) was added to each sample and the homogenization repeated. Homogenized samples in 2.5% LiDS were heated to 95 C for 5 minutes then filtered through spin columns (Pierce #89868) to produce a clear lysate solution. An aliquot of each filtered lysate was taken for protein analysis then the remainder of each sample was buffered with 20 mM Tetraethyammonium bicarbonate (TEAB, Honeywell Fluka cat# 60-044-974) then reduced with 5 mM dithiothreitol (DTT) and alkylated with 15 mM iodoacetamide (IAA) with excel IAA quenched after a 30 minute incubation by addition of a second aliquot of 5 mM DTT. Samples were acidified by addition of 20 ul of 12% phosphoric acid then proteins precipitated by addition of 1 ml of 90% MeOH in 100 mM TEAB. Pellets from the precipitation were washed with 0.5 ml of 80% MeOH in 10 mM TEAB. Precipitates were dried on the bench then resuspended in 150ul of 100mM NaCl, 1mMCacl, 40mM TEAB and 0.5% deoxycholate (DOC). Trypsin (Promega, V5113) was added to each sample, 1ug per sample, and then incubated overnight at 37 C to complete the digestion.

### Mass spectrometry analysis

LC-MS/MS analysis was performed using a Thermo scientific Vanquish-Neo chromatography system with an Acclaim PepMap 100 trap column (100 µm × 2 cm, C18, 5 µm, 100Å), and Thermo Scientific Easy-Spray PepMap RSLC C18 75 um x 25 cm column. A gradient starting at 6% of a solution of 80% Acetonitrile with 0.1% Formic acid and finishing at 42% acetonitrile 60 minutes later was used for all samples. Data independent analysis was performed on an Orbitrap Eclipse MS system. MS1 spectra were acquired at 120,000 resolution in the 350 to 1650 Da mass range with an AGC of 3e6. MS2 spectra were acquired in the Orbitrap and collected at 15,000 resolution with a 200 to 1600 Da window. Fragmentation for MS2 spectra was performed by HCD with a collision energy of 30, a maximum injection time of 120 msec, and an AGC target of 3e6. Variable windows of 23 to 54 Da were used for 21 windows between 377 and 972 Da. A window of 71 Daltons was centered on 1034.5, a window of 194 was centered on 1133.5, and a window of 453 was centered on 1423.5 to complete the acquisition. Mass spectrometry data were processed by Spectronaut 18.6 (Biognosis), using BGS factory settings normalization and Mouse Uniprot FASTA database (downloaded March 30, 2021, 17,035 entries). The search parameters included trypsin with up to two missed cleavage. Variable modification by oxidation of M, and of protein N-termini by Acetylation, Met Loss or both Acetylation and Met loss. Carbamidomethylation of cysteine was a fixed modification. For the entire data set, false discovery rate (FDR) was calculated using a cut-off of 1% for identification of precursors and 1% for identifications of peptides. Proteins with more than 50% missing values across the 12 samples were excluded, and missing values in the remaining dataset were imputed using a random forest-based imputation method. Protein abundance levels were normalized by the median value of each sample, followed by log2 transformation to stabilize variance. Quantitative protein profiling was evaluated using Principal Component Analysis (PCA) and Partial Least Squares-Discriminant Analysis (PLS-DA) to assess group discrimination. PCA and PLS-DA score plots are shown in **Figure S2C, D.** For differential analysis, two-sample t-tests were performed, followed by an FDR correction using the Benjamini-Hochberg method. Differentially abundant proteins were identified based on an unadjusted p-value threshold of 0.05 and a fold-change cutoff of 1.1. Pathway analysis was conducted using iPathwayGuide (Advaita Bioinformatics, Ann Arbor, MI, USA).

### Statistical analysis

For statistical significance, data were analyzed using Statistica software (version 14.0.015). For the GTT, ITT, and energy expenditure, a repeated measures analysis of variances (ANOVA) was used. For all ANOVA analyses, a Newman-Keuls post hoc analysis followed. For any data sets where only 2 groups of the predictor value were being compared, a two-tail unpaired Student’s *t*-test was performed. For all statistical analyses, a 5% level of significance was set.

## Results

### 1. Long-term benzene exposure at occupational levels induces metabolic dysfunction in adult male mice

To model chronic exposure to benzene at sub-regulatory occupational levels, we exposed adult male mice to 0.9 ppm benzene for 8 hours per day over 9 weeks (**Fig. 1A**). Following exposure, urinary analysis indicated elevated levels of the benzene metabolite trans, trans-muconic acid (tt-MA) in urine of exposed males, though this increase did not reach statistical significance (**Fig. S1**). Throughout the exposure, no changes in body weight or body composition were observed (**Fig. 1B** and **C**). However, by week 5, benzene-exposed mice exhibited glucose intolerance, as shown by impaired glucose tolerance test (GTT) (**Fig. 1D, E**). This was further increased by week 8, with benzene-exposed mice developing hyperglycemia (**Fig. 1F, G**). Furthermore, insulin tolerance test (ITT) demonstrated a marked reduction in insulin sensitivity in benzene-exposed male mice (**Fig. 1H**), with elevated fasting insulin levels and increased Homeostatic Model Assessment for Insulin Resistance (HOMA-IR) scores following 8 weeks of exposure (**Fig. 1I, J**). These findings establish that long-term exposure to benzene at levels below occupational threshold is sufficient to induce significant metabolic dysfunction, characterized by glucose intolerance and insulin resistance.

### 2. Long-term occupational benzene exposure alters energy expenditure

To further understand the effects of occupational benzene exposure on whole-body energy homeostasis, we examined parameters related to energy expenditure. Unexpectedly, prolonged occupational benzene exposure led to significantly increased oxygen consumption (O₂), carbon dioxide production (CO₂), and energy expenditure (EE) during the dark cycle (**Fig. 2A-C**), with no changes in total activity and food consumption (**Fig. 2D-E**). In the context of observed insulin resistance and glucose intolerance, the elevated EE might represent a form of metabolic stress driven by systemic inflammation, particularly in insulin-sensitive tissues [18].

**Figure 2.**
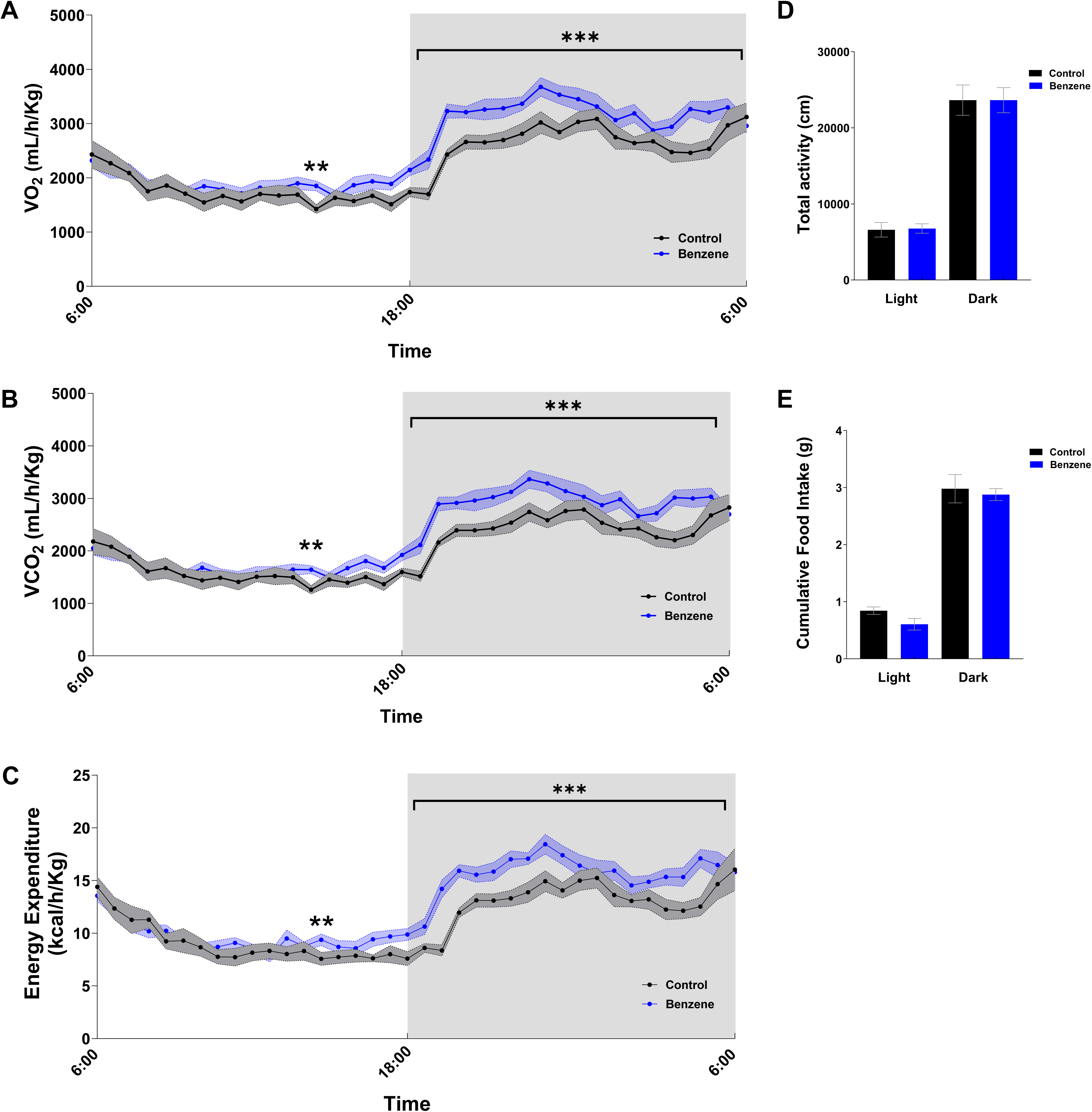
Prolonged occupational benzene exposure impairs energy homeostasis. Energy homeostasis parameters measured during the light and dark cycle over 48h (A) Oxygen consumption (VO_2_) [ml/h/kg], (B) Carbon dioxide production (VCO_2_) [ml/h/kg], (C) Energy expenditure [kcal/h/;kg], (D) Locomotor activity, and (E) Cumulative food intake (g). All data are shown as the mean ± SEM. Repeated measure ANOVA followed by Newman-Keuls post-hoc test, or student’s t-test. (*,p<0.05, **, p<0.01, ***,p<0.001; n=6/group).

### 3. Benzene exposure alters the transcriptome of insulin-sensitive tissues

To assess the impact of occupational benzene exposure on peripheral insulin-sensitive organs, we performed bulk RNA-seq on liver, skeletal muscle, and adipose tissue from benzene-exposed male mice.

### a. Liver

In the liver, we identified 101 downregulated and 145 upregulated genes (DEGs) in response to long-term exposure to benzene. There was strong downregulation of genes related to gluconeogenesis and lipid metabolism, such as *Pdk4* and *Asap2*, as well as *Lgals1*, which is involved in modulating inflammatory responses and immune homeostasis [19–21]. Among the top upregulated genes, we identified *Bhlhe41, Fpr1, Fpr2,* and *Cyp2b9*, indicating hepatic stress adaptation, metabolic regulation, innate immune responses, and detoxification processes triggered by benzene exposure [22–24] (**Fig. 3A**). Pathway enrichment analysis revealed the top enriched Biological Process (BP), Cellular Component (CC), and Molecular Function (MF) terms, mostly related to pathways critical for immune function and stress responses. These included ‘antigen processing and presentation of peptide antigen via MHC class II’, ‘Fc receptor signaling pathway’, ‘response to interferon-beta’, ‘Interleukin-1 beta production’, ‘inflammatory response’, ‘response to stress’, ‘MAP kinase phosphatase activity’, and ‘phospholipase A2 activity’, reflecting immune activation in liver tissue, modulation of cytokine production, stress signaling, and enzymatic regulation of inflammatory mediators in response to benzene exposure (**Fig. 3B-C**). Further, KEGG enrichment analysis demonstrated pathways related to immune activation, including ‘FCGR activation’, ‘macrophage markers’, ‘Type 1 Diabetes mellitus’, ‘PI3K/Akt signaling’, ‘phagosome’, and ‘NOD-like receptor signaling’, suggesting the involvement of detoxification processes and metabolic dysfunction in the liver (**Fig. 3D**). **Figures 3E** and **3F** present heatmaps for some of the top immune and metabolic genes activated by benzene exposure in the liver. Thus, even low-level benzene exposure was sufficient to drive liver inflammation, triggering insulin resistance and metabolic dysfunction.

**Figure 3.**
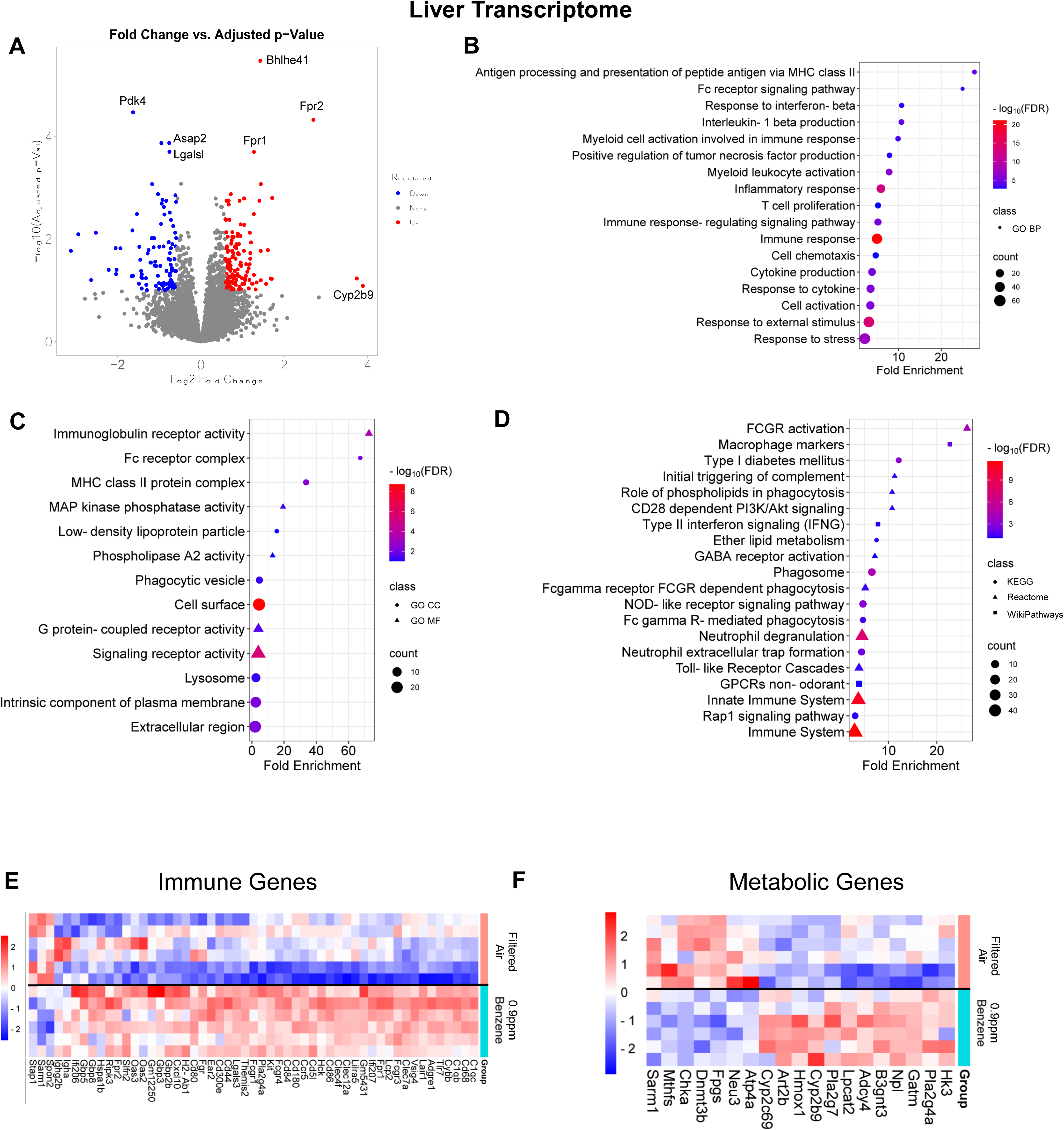
Hepatic transcriptome analysis following benzene exposure. (A) Volcano plot of top DEGs (FDR<0.1, FC >1.5). (B) Pathway enrichment for Biological process (BP). (C) Pathway enrichment for Cellular component (CC) and Molecular function (MF). (D) Pathway enrichment for KEGG, Reactome, and WikiPathways. (E) Top DEGs from BP immune (E) and (F) KEGG metabolic pathways. Pathway analyses, FDR<0.05; n=3-6/group.

### b. Muscle

In skeletal muscle, our transcriptomic analysis identified 378 upregulated and 251 downregulated DEGs compared to filtered-air control mice. Among the top downregulated genes, we identified genes involved in muscle stress and damage such as *Grem2, Enox2*, and *Golm1*, while *Lrtm2, Pitx2, Ankrd33b, Mybpc1,* and *Egln3* were upregulated (**Fig. 4A**) [25–30]. Top GO BP, CC, and MF terms included ‘inositol phosphate dephosphorylation’, ‘muscle cell development’, ‘muscle contraction’, ‘inositol phosphate activity’, ‘CHOP-ATF3 complex’, and ‘cytoskeletal motor activity’, among others (**Fig. 4B-C**). Interestingly, KEGG, Reactome, and WikiPathways revealed enrichment in ‘HIF-1 signaling pathway’, ‘Wnt signaling pathway’, ‘alpha-6 beta-4 integrin signaling pathway’ and ‘hypoxia dependent proliferation of myoblasts’ suggesting a hypoxia-like stress responses in muscle tissue (**Fig. 4D**). Regardless, some top upregulated genes included those with known roles in disrupted insulin signaling, metabolic regulation, and oxidative stress response, such as *Pde10a, B3galt1, Aldh1a1, Sgms2, Adh1,* and *Galnt15* [31–34]. On the other hand, some key downregulated genes, such as *Gadl1, Amd1, Cyp26b1, Pde7a, Shmt1, B3galnt2, Tecr,* and *Adi1*, are integral to metabolic homeostasis and cellular maintenance [35–40] (**Fig. 4E** and **F**), thus suggesting compromised cellular maintenance and metabolic adaptation.

**Figure 4.**
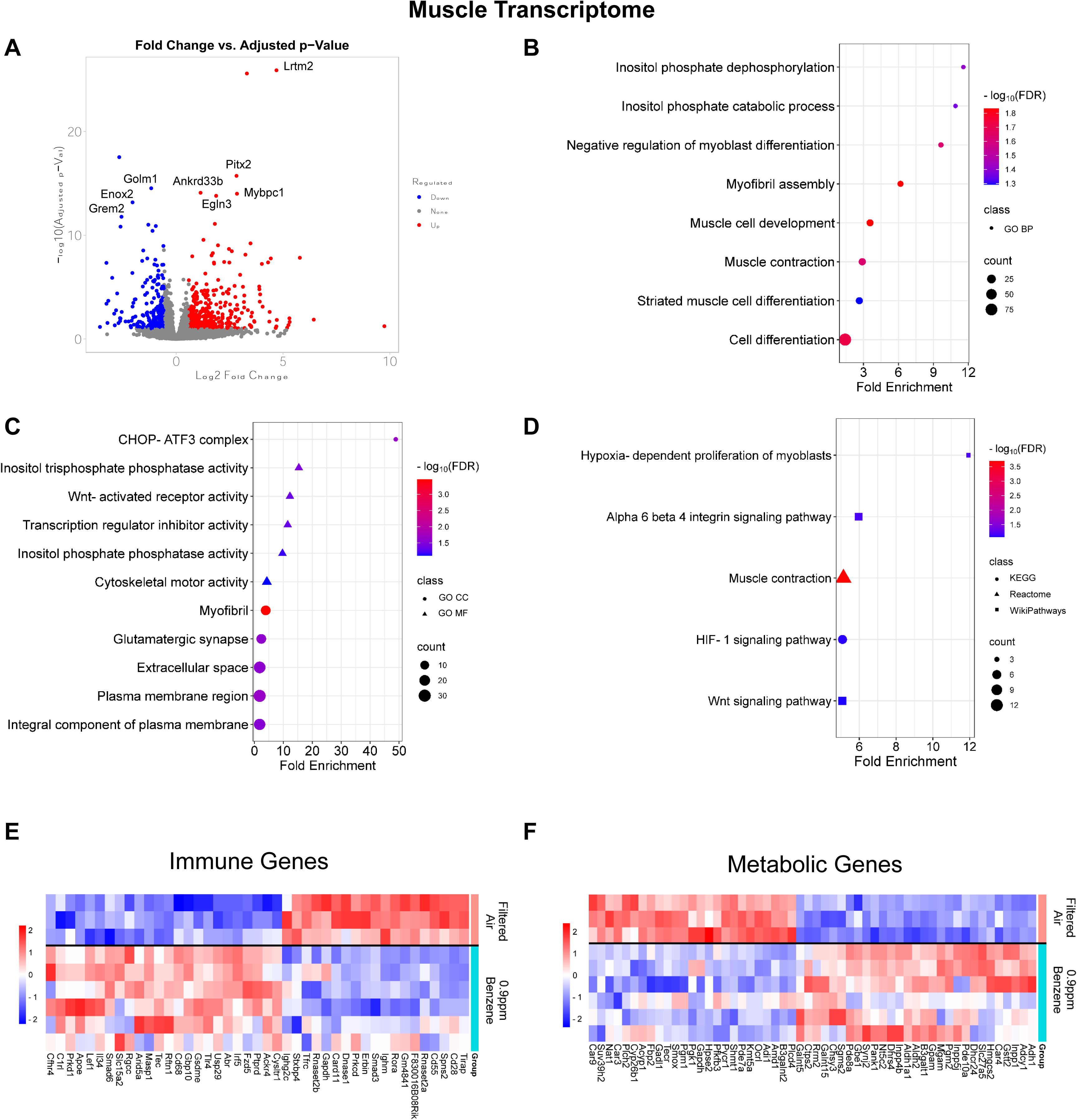
Muscle transcriptome analysis following benzene exposure. (A) Volcano plot of the top DEGs (FDR <0.1, FC >1.5). (B) Pathway enrichment for Biological process (BP). (C) Pathway enrichment for Cellular component (CC) and Molecular function (MF). (D) Pathway enrichment for KEGG, Reactome, and WikiPathways. Top DEGs from BP immune (E) and (F) KEGG metabolic pathways. Pathway analyses, FDR<0.05; n=3-6/group.

### c. Adipose tissue

No significant DEGs were identified in adipose tissue following exposure, indicating a lack of a direct transcriptional response at this dose (#PRJNA1194603).

### 4. Benzene exposure alters hepatic proteome

Since the most apparent metabolic and inflammatory changes in response to benzene exposure were observed in the liver, we next examined the hepatic proteome to further explore these changes at the protein level. We identified 236 proteins with significant abundance difference between groups with 100 proteins having higher abundance in benzene-exposed and 136 proteins having higher abundance in the control group (**Fig. 5A**). In support with our transcriptomics data, pathway enrichment analysis of the hepatic proteome identified significant involvement in ‘mitophagy’, ‘oxidative phosphorylation’ and ‘NOD-like receptor signaling’ indicating the disrupted liver metabolic homeostasis and immune regulation [41–43], that can contribute to the pro-inflammatory environment. Furthermore, GO analysis revealed enrichment in pathways related to mitochondrial fusion, localization, and cellular catabolic processes suggesting that mitochondrial function in liver is directly impacted by the benzene exposure at occupational levels (**Fig. 5B**). In support, we identified significant changes in key immune and metabolic proteins linked to mitochondrial function, including the upregulation of immune proteins such as Crk, C1qb, and Gsdmd [44, 45], and the downregulation of metabolic proteins such as Ndufb8, Fmo5, and Atp5o [46, 47], suggesting impaired mitochondrial energy production (**Fig. 5C-F**). While there was minimal overlap between our transcriptomic and proteomic results at the individual gene level (limited to *C1qb*, *Hspa1b*, and *Mrc1*), the upregulation of these proteins, together with pathways effects, emphasizes the activated immune response to mitochondrial stress, oxidative damage, and inflammation [45, 48]. Together, these changes suggest mitochondrial dysfunction as a central driver of the pro-inflammatory state and metabolic imbalance in benzene-exposed liver.

**Figure 5.**
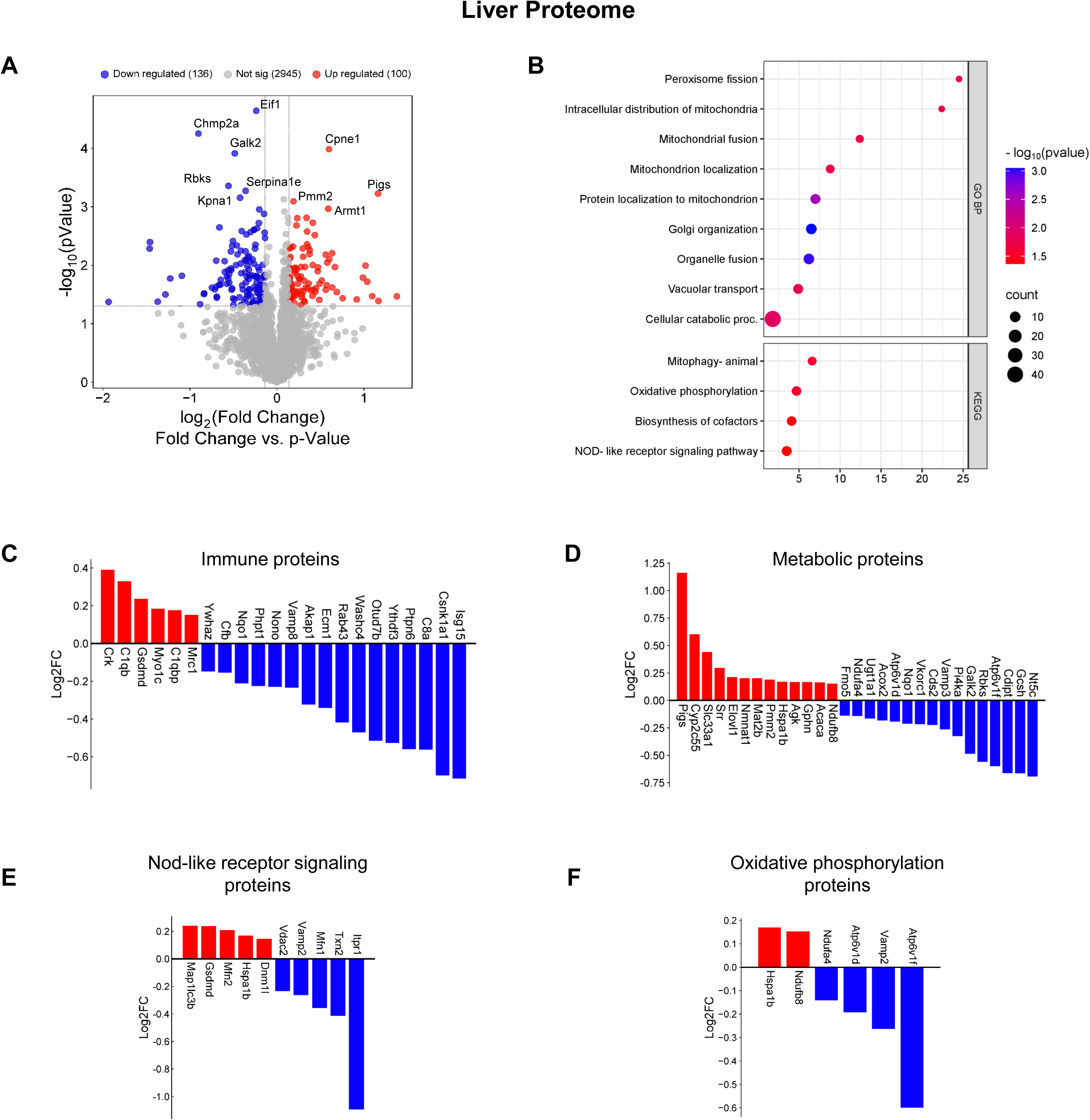
Hepatic proteome analysis following benzene exposure. (A) Volcano plot of the top significantly abundant proteins (p value <0.05, FC <1.1). (B) Pathway enrichment for Biological process and KEGG pathway. (C) Immune, (D) metabolic, (E) Nod-like receptor signaling, and (F) Oxidative phosphorylation differentially abundant proteins from KEGG pathway enrichment. (G) Overlap of genomics and proteomics data analysis. All pathway analyses used FDR <0.1; n=6/group.

### 5. Comparative analysis of hepatic transcriptome to occupational and smoking levels of benzene

Our previous studies demonstrated that benzene exposure at smoking levels (50 ppm) induces hyperglycemia and hyperinsulinemia after 4 weeks of exposure [12]. These metabolic disruptions were associated with increased expression of hepatic inflammatory genes and impairments in blood lipids [12]. To identify both shared and unique hepatic transcriptomic signatures between benzene exposure levels contributing to whole-body metabolic imbalance, we compared the hepatic expression profiles of mice exposed to high (50 ppm) and low (0.9 ppm) doses of benzene. Volcano plot demonstrates upregulated (red) and downregulated (blue) DEGs in liver following exposure to 50 ppm benzene (**Fig. 6A**). Some upregulated genes, including *Per3, Slc16a12, Wee1*, and *Dbp*, are associated with circadian rhythm, cell cycle regulation, and metabolic stress. Pathway enrichment analysis demonstrated that benzene exposure at smoking levels significantly impacted pathways related to SREBP signaling, circadian rhythm regulation, arachidonic acid metabolism, and ER-nucleus signaling. In support with our previous work [12], metabolic pathways such as cholesterol and lipid metabolism, steroid hormone biosynthesis, retinol metabolism, and PPAR signaling also showed significant enrichment (**Fig. 6B-D**). Importantly, pathways involving cytochrome P450 activity, such as oxidation by cytochrome P450, arachidonic acid metabolism, and steroid hormone metabolism, were among the most enriched KEGG pathways (**Fig. 6D**), indicating the liver metabolic and detoxification response to benzene exposure. Among the shared genes across both exposure levels, key metabolic and stress-response genes were identified. Specifically, some of these genes play critical roles in detoxification and xenobiotic metabolism (*Akr1c19* and *Cyp2b9*) [49, 50], circadian rhythms (*Dbp* and *Ciart*) [51], oxidative stress regulation and mitochondrial integrity (*Scara5* and *Wee1*) [52], and insulin signaling and inflammatory responses (*Usp2*) [53](**Fig. 6E**). To explore shared transcriptomic signatures, we performed a Cytoscape pathway analysis on all statistically significant DEGs. The analysis revealed that both exposure levels shared enrichment in pathways related to cellular aromatic compound metabolic processes, which likely reflect benzene metabolism (**Fig. 6F**). Additional shared pathways included ‘cellular metabolic processes’, ‘cellular communication’, and various organelle-related functions, such as ‘transport and mitochondrial membrane organization’ (**Fig. 6F**). These shared pathways suggest that, irrespective of dose, benzene exposure disrupts critical cellular processes, leading to altered cellular function and systemic metabolic imbalance.

**Figure 6.**
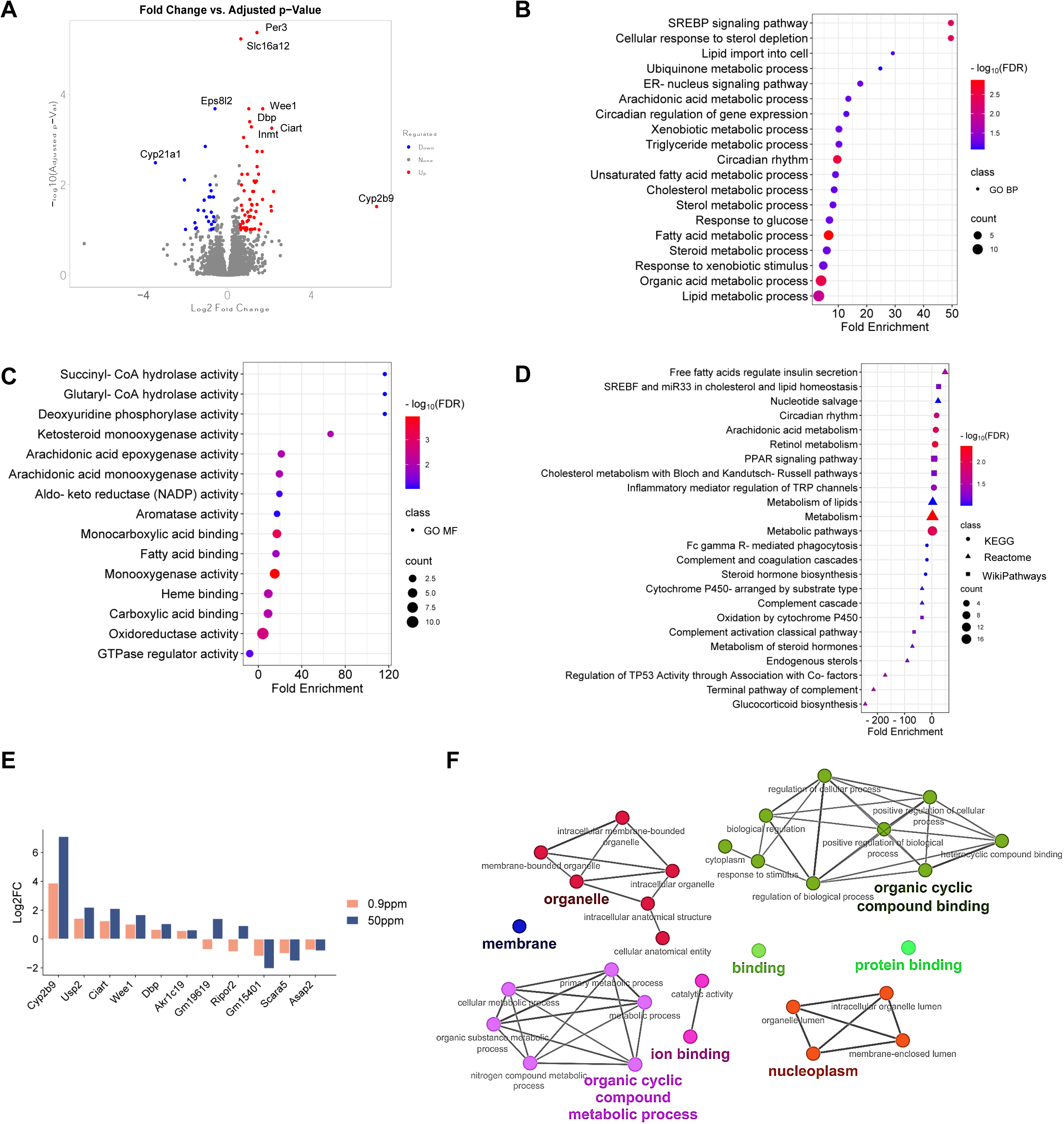
Comparative analysis of hepatic transcriptome to occupational and smoking levels of benzene. (A) Volcano plot of top DEGs in liver (50 ppm, FDR <0.1, FC >1.5). Pathway enrichment for (B) Biological process and (C) molecular function (50 ppm, FDR <0.05). (D) Pathway enrichment of KEGG, Reactome, and WikiPathways (50 ppm, FDR <0.05). (E) Shared DEGs in liver of 50 ppm and 0.9 ppm benzene exposure. (F) Interaction network of shared GO terms between 50 ppm and 0.9 ppm levels (p value <0.05; n=3-6/group).

## Discussion

Our study is the first to investigate the impact of prolonged, low-dose benzene exposure at occupationally relevant dose on whole-body metabolism. We demonstrate that chronic benzene exposure significantly disrupts glucose homeostasis, induces insulin resistance, alters energy expenditure and reshapes the transcriptional profile of liver and muscle, two key metabolic insulin-sensitive tissues. Importantly, these metabolic disturbances occurred in the absence of changes in body weight, body composition, or transcriptional changes in adipose tissue, suggesting a direct effect of the exposure on metabolic regulation rather than secondary effect of weight gain. Using a multiomics approach, we identified the liver as a primary site of benzene-induced metabolic disruption, driven by immune activation and mitochondrial stress responses. Importantly, these pathways were conserved across smoking-related and occupational levels. Our findings emphasize the potential metabolic health risks associated with occupational benzene exposure, even at sub-regulatory levels.

Mice exposed to benzene at occupational levels developed glucose intolerance, hyperglycemia, and insulin resistance with worsening over time. Interestingly, the observed metabolic imbalance was accompanied by increased energy expenditure, with no changes in food intake, or locomotor activity in these animals. The elevated energy expenditure likely reflects underlying metabolic stress caused by systemic inflammation and mitochondrial dysfunction in liver [18, 54]. Indeed, chronic low-grade inflammation and impaired mitochondrial energy production have been shown to elevate systemic energy expenditure by as much as 10% [55], [56]. In support, our liver multiomics analysis revealed significant changes in mitochondrial pathways, such as oxidative phosphorylation and mitophagy, indicating the role of mitochondrial stress in mediating benzene-induced metabolic disruption. Upregulation of hepatic genes involved in stress and immune responses, such as *Cyp2b9, Fpr1*, and *Fpr2,* and downregulation of genes like *Pdk4*, and *Lgals1* [19–21] further demonstrate the impact of benzene exposure on hepatic glucose and lipid metabolism. The altered abundance of immune-related proteins, such as C1qb and Mrc1 [45, 48], and metabolic proteins like Ndufb8 and Fmo5 reflects impaired mitochondrial energy production and detoxification [46, 47]. Collectively these effects may contribute to systemic insulin resistance and glucose intolerance.

In our previous study, using mice exposed to benzene at smoking levels (50 ppm), we demonstrated similar metabolic impairments, including hyperglycemia, hyperinsulinemia, and insulin resistance [4, 12]. This high-dose exposure induced marked inflammatory and immune activation pathways, including SREBP signaling, NOD-like receptor signaling, and ER-nucleus signaling, reflecting a more severe hepatic response as compared to the lower dose. Regardless, we reveal the conserved transcriptomic signatures in liver between the two exposure levels, such as upregulation of key stress-response genes (*Akr1c19, Dbp, Wee1, and Cyp2b9*) [57–59] and enrichment of pathways involved in xenobiotic metabolism, circadian rhythm regulation, and cytochrome P450 activity. These shared responses suggest that mitochondrial dysfunction and oxidative stress are central mechanisms driving benzene-induced metabolic dysfunction, regardless of dose.

Skeletal muscle exhibited a modest transcriptomic response, with changes in a subset of genes related to stress adaptation, metabolic regulation, and cellular maintenance. For example, *Egln3* was upregulated, suggesting compensatory responses to mitigate oxidative stress, while downregulated genes like *Grem2* and *Shmt1* suggest impairments in cellular maintenance and one-carbon metabolism, critical for muscle homeostasis [25, 30]. However, the absence of enrichment in metabolic pathways in skeletal muscle suggests a secondary role in benzene-induced metabolic dysfunction. Adipose tissue, in contrast, showed no significant transcriptomic changes in response to benzene exposure, indicating a lack of direct impact on this tissue at the transcriptional level. Previous studies have suggested that adipose tissue can buffer systemic toxic exposure by storing lipophilic chemicals, reducing their bioavailability and downstream impact on metabolism [60]. Both liver and muscle are highly metabolically active and dependent on mitochondrial oxidative phosphorylation, while adipose tissue has a lower mitochondrial density and thus less involved in xenobiotic metabolism, possibly protecting it from the immediate effects of benzene exposure [61] [62].

Our findings raise critical concerns about the levels of current exposure standards. The observed glucose intolerance, insulin resistance, and hepatic mitochondrial dysfunction highlight benzene potential to disrupt systemic metabolic pathways, even at levels below regulatory threshold. Importantly, our findings are consistent with epidemiological studies linking benzene exposure to increased risks of metabolic syndrome and T2DM in exposed populations [4]. Given the central role of the liver in benzene metabolism, the observed hepatic inflammation and mitochondrial dysfunction may amplify these risks by increasing systemic inflammation and oxidative stress.

### Study Limitations

In current study we exclusively focused on male mice, limiting our understanding of potential sex-specific responses to exposure. In our previous studies, we did not detect any significant metabolic abnormalities in female mice exposed to a higher dose benzene for up to 4 weeks [4, 12]. However, sex-specific differences are still possible on transcriptional level, and will be addressed in the future. Furthermore, longer-term studies are necessary to assess cumulative effects and progressive damage that may be noticeable over time in other organs, including adipose tissue.

Together, our findings reveal the systemic impact of OSHA approved benzene exposure levels on peripheral metabolism. We demonstrate that even low-dose exposure, previously considered safe, is sufficient to induce significant metabolic disruptions, emphasizing the potential health risks associated with occupational benzene exposure. These data highlight the need for stricter regulatory standards to protect exposed populations from chronic metabolic and inflammatory diseases.

## Acknowledgments

This project was supported by CURES Center Grant (P30ES036084), NIEHS R01ES033171, and CLEAR P42ES030991 for MS. NIEHS R56-ES034765 further supported MS, AL, AB, and SS. LK was further supported by the NIH T32GM142519 and LS was supported by NIH 5T32HL120822-09. We acknowledge the support of the Natural Sciences and Engineering Research Council of Canada (NSERC), #578118, as well as SNAP provided by the WSU Pharmacology department to RB. We acknowledge the assistance of the WSU Proteomics Core that is supported through NIH P30 ES036084, P30 CA022453 and S10 OD030484. The author(s) thank the Van Andel Institute Genomics Core Facility (RRID:SCR_022913) for providing RNA-sequencing facilities and services. CHL was supported by Centers for Disease Control and the National Institute on Drug Abuse of the National Institutes of Health under award number U01DA053893-01. We would also like to thank the Center for Agroforestry at the University of Missouri, USDA/ARS Dale Bumpers Small Farm Research Center under agreement number 58-6020-6-001 from the USDA Agricultural Research Service for supporting part of this research. The Biostatistics and Bioinformatics Core is supported, in part, by NIH P30 CA022453 to the Karmanos Cancer Institute at WSU. The Wayne State Genome Sciences Core is further supported by CURES P30 ES036084. Figure graphics were created with BioRender.com.

## Data Availability Statement

The data that support the findings of this study are available from the corresponding author upon reasonable request.

## Abbreviations

VOCs: Volatile organic compounds

T2DM: Type 2 Diabetes Mellitus

CYP: Cytochrome P450

ppm: Parts per million

## Supplementary Material

**Figure S1.** Urinary metabolite tt-MA. (A). Urinary tt-MA, normalized to creatinine levels. Data are shown as the mean ± SEM (n=5-9/group).

**Figure S2.** Principle Component Analysis (PCA) for transcriptomic and proteomic analyses. (A) PCA score plot of 0.9 ppm and 50 ppm liver RNA samples. (B) PCA score plot of 0.9ppm benzene muscle RNA versus filtered air controls. (C) PCA score plot of 0.9 ppm benzene liver protein versus filtered air controls. (D) Partial Least Squares Discriminant Analysis (PLS-DA) score plot for 0.9 ppm benzene liver protein versus filtered air controls. The circles represent 95% confidence ellipses.

## Notes

https://www.ncbi.nlm.nih.gov/bioproject/1194550

https://www.ncbi.nlm.nih.gov/bioproject/1194603

### Competing Interest Statement

The authors have declared no competing interest.

