## Supplementary Figures for "Integrative multiomics analysis of metabolic dysregulation induced by occupational benzene exposure in mice"

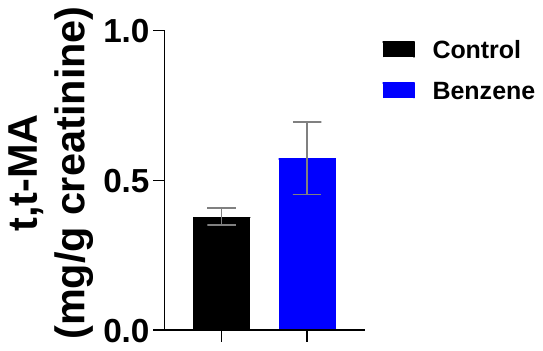

**Figure S1.** **Urinary metabolite tt-MA**. (A). Urinary tt-MA, normalized to creatinine levels. Data are shown as the mean ± SEM (n=5-9/group).

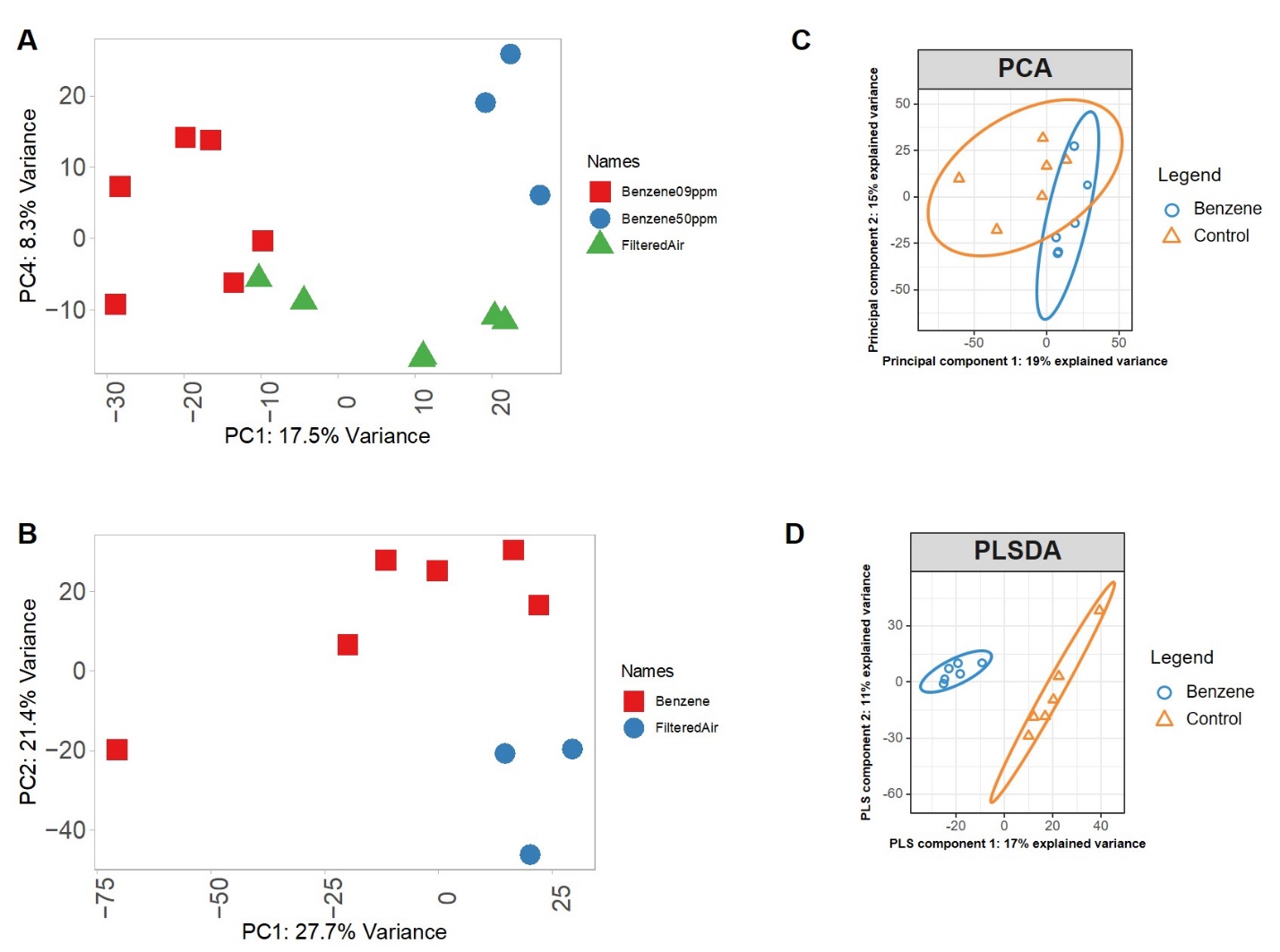

**Figure S2. Principle Component Analysis (PCA) for transcriptomic and proteomic analyses.** (A) PCA score plot of 0.9 ppm and 50 ppm liver RNA samples. (B) PCA score plot of 0.9ppm benzene muscle RNA versus filtered air controls. (C) PCA score plot of 0.9 ppm benzene liver protein versus filtered air controls. (D) Partial Least Squares Discriminant Analysis (PLS-DA) score plot for 0.9 ppm benzene liver protein versus filtered air controls. The circles represent 95% confidence ellipses.
